# 7 Tesla MRI of the *ex vivo* human brain at 100 micron resolution

**DOI:** 10.1101/649822

**Authors:** Brian L. Edlow, Azma Mareyam, Andreas Horn, Jonathan R. Polimeni, Thomas Witzel, M. Dylan Tisdall, Jean Augustinack, Jason P. Stockmann, Bram R. Diamond, Allison Stevens, Lee S. Tirrell, Rebecca D. Folkerth, Lawrence L. Wald, Bruce Fischl, Andre van der Kouwe

**Author notes:** co-senior authors (Bruce Fischl, Andre van der Kouwe). corresponding author: Brian Edlow.

## Abstract

We present an ultra-high resolution MRI dataset of an *ex vivo* human brain specimen. The brain specimen was donated by a 58-year-old woman who had no history of neurological disease and died of non-neurological causes. After fixation in 10% formalin, the specimen was imaged on a 7 Tesla MRI scanner at 100 μm isotropic resolution using a custom-built 31-channel receive array coil. Single-echo multi-flip Fast Low-Angle SHot (FLASH) data were acquired over 100 hours of scan time (25 hours per flip angle), allowing derivation of a T1 parameter map and synthesized FLASH volumes. This dataset provides an unprecedented view of the three-dimensional neuroanatomy of the human brain. To optimize the utility of this resource, we warped the dataset into standard stereotactic space. We now distribute the dataset in both native space and stereotactic space to the academic community via multiple platforms. We envision that this dataset will have a broad range of investigational, educational, and clinical applications that will advance understanding of human brain anatomy in health and disease.

|  |  |
| --- | --- |
| **Design Type(s)** | Single measure design |
| **Measurement Type(s)** | Nuclear magnetic resonance assay |
| **Technology Type(s)** | 7 Tesla MRI scanner |
| **Factor Type(s)** |  |
| **Sample Characteristic(s)** | Homo sapiens • brain |

## Background & Summary

Postmortem *ex vivo* MRI provides significant advantages over *in vivo* MRI for visualizing the microstructural neuroanatomy of the human brain. Whereas *in vivo* MRI acquisitions are constrained by time (i.e. ~hours) and affected by motion, *ex vivo* MRI can be performed without time constraints (i.e. ~days) and without cardiorespiratory or head motion. The resultant advantages for characterizing neuroanatomy at microscale are particularly important for identifying cortical layers and subcortical nuclei^1–5^, which are difficult to visualize even in the highest-resolution *in vivo* MRI datasets^6,7^. *Ex vivo* MRI also provides advantages over histological methods that are associated with distortion and tearing of human brain tissue during fixation, embedding, and slide-mounting.

As the field of *ex vivo* MRI has developed over the past two decades, several laboratories have focused on imaging blocks of tissue from human brain specimens using small-bore scanners^2,8^ and specialized receive coils^9–11^. This approach allows for spatial resolutions of up to 35–75 microns for analyses of specific neuroanatomic regions^9,11–13^. However, ultra-high resolution imaging of whole human brain specimens at high magnetic field strengths has been far more challenging, due to the need for multi-channel receive coils and large-bore clinical scanners that can accommodate a whole-brain specimen. Whole-brain imaging is required to observe neuroanatomic relationships across distant brain regions, as well as to provide a complete view of human neuroanatomy in standard stereotactic space.

Here, we report the results of a multidisciplinary effort to image a whole human brain specimen *ex vivo* at an unprecedented spatial resolution of 100 μm isotropic. Central to this effort was the construction of an integrated system consisting of a custom-built 31-channel receive array coil and volume transmit coil, which was designed to accommodate and tightly enclose an *ex vivo* human brain^14^. The scans were performed on a 7 Tesla whole-body human MRI scanner using four single-echo spoiled gradient-recalled echo (SPGR/GRE) or Fast Low-Angle SHot (FLASH) sequences. We used varying flip-angles (FA15°, FA20°, FA25°, FA30°) to generate multiple synthesized volumes, each of which provides a different tissue contrast. The scans, performed over ~100 hours (~25 hours per FA), generated an ~8 TB dataset (~2 TB per flip angle) that required custom-built computational tools for offline MRI reconstruction and creation of the synthesized volumes. Offline MRI reconstruction considerably reduces the data amount. We release the resulting FA25° acquisition, as well as the synthesized FLASH25 volume here, both in native space and coregistered to standard stereotactic space, for use by the academic community. We envision a broad range of investigational, educational, and clinical applications for this dataset that have the potential to advance understanding of human brain anatomy in health and disease.

## Methods

### Specimen acquisition and processing

A 58-year-old woman with a history of lymphoma and stem cell transplantation, but no history of neurological or psychiatric disease, died in a medical intensive care unit. She was initially admitted to the hospital for fevers, chills, and fatigue, and then was transferred to the intensive care unit for hypoxic respiratory failure requiring mechanical ventilation. Her hospital course was also notable for a deep venous thromboses and a pulmonary embolism. The cause of her death on hospital day 15 was determined to be hypoxic respiratory failure due to viral pneumonia. At the time of her death, her surrogate decision-maker provided written informed consent for a clinical autopsy and for donation of her brain for research, as part of a protocol approved by our Institutional Review Board.

At autopsy, her fresh brain weighed 1,210 grams (normal range = 1,200 to 1,500 grams). The brain was fixed in 10% formalin 14 hours after death. Gross examination revealed a normal brain (Fig. 1), without evidence of mass lesions or cerebrovascular disease. To ensure adequate fixation and prevent specimen flattening (which can prevent specimens from fitting into custom *ex vivo* MRI coils), we followed a series of standard specimen processing procedures, as previously described^15^.

**Figure 1.**
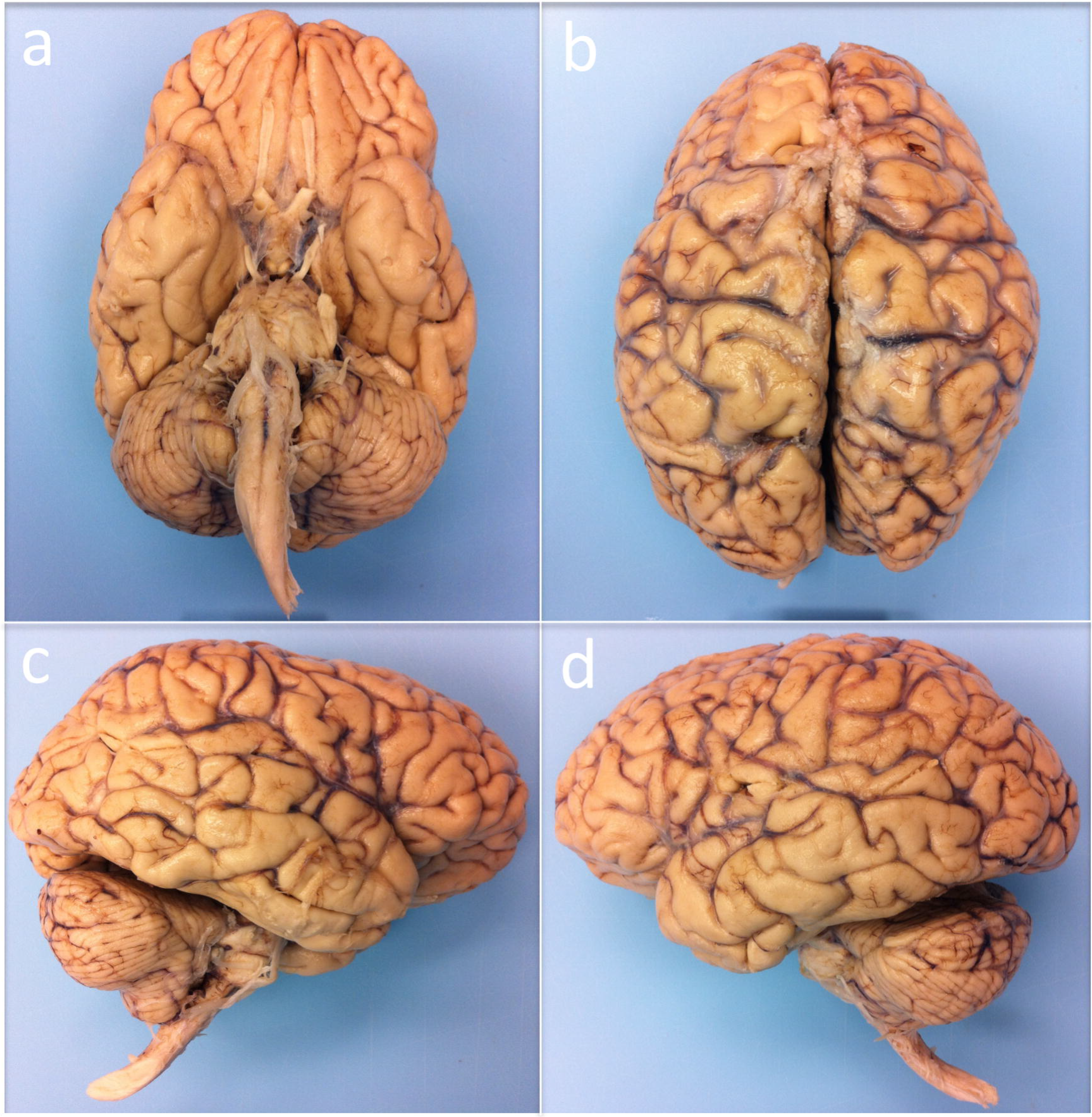
Human brain specimen. The human brain specimen that underwent *ex vivo* MRI is shown from inferior (**a**), superior (**b**), right lateral (**c**) and left lateral (**d**) perspectives. Gross pathological examination of the brain was normal.

### Specimen preparation for scanning

After remaining in fixative for 35 months, the brain specimen was transferred to Fomblin Y LVAC 06/6 (perfluoropolyether, Solvay Specialty Polymers USA, LLC, West Deptford, NJ), which is invisible to MR and reduces magnetic susceptibility artifacts. The specimen, immersed in Fomblin, was then secured inside a custom-built, air-tight brain holder made of rugged urethane^16^. The brain holder contains degassing ports for removal of air bubbles, which further reduces magnetic susceptibility artifacts.

### Construction of a receive array coil and transmit volume coil for *ex vivo* imaging of the whole human brain

We built a receive coil apparatus consisting of a 31-channel surface coil loop array with two halves. The apparatus was fabricated using a 3D printer of slightly larger dimensions than the brain holder, which slides inside the single-channel birdcage volume transmit coil (Fig. 2). The brain holder is an oblate spheroid (16 × 19 cm) that conforms to the shape of a whole brain (cerebral hemispheres + cerebellum + brainstem)^16^ (Fig. 2d). It is made of two separate halves that can be sealed together with a silicone gasket after packing the brain inside. This holder must also withstand the degassing process when under vacuum pressure. Degassing is performed in three steps: 1) introducing vacuum suction into the container with the brain inside, which allows the bubbles to expand under decreased pressure and exit tissue cavities; 2) opening the valve to fill the holder with fomblin and then sealing off the fill valve; and 3) continuation of vacuum suction with low-amplitude vibration of the holder for 2-6 hours. The vibration facilitates the removal of bubbles from tissue cavities. All three steps are performed inside a fume hood.

**Figure 2.**
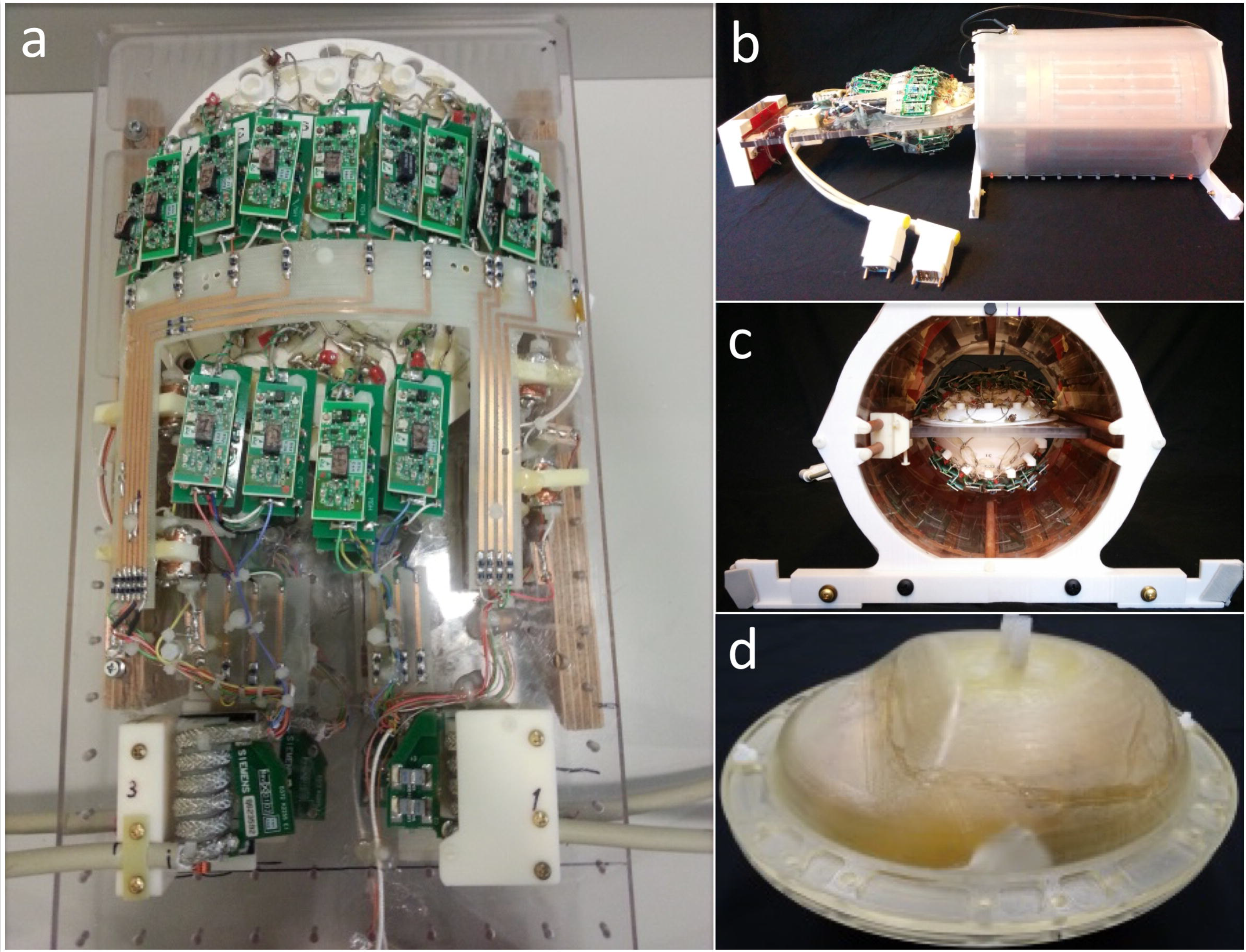
Receive array coil and transmit volume coil for *ex vivo* imaging of the whole human brain. (**a**) The 31-channel receive array has 15 elements on the top half (with a diameter of 5.5 cm) and 16 on the bottom half (with a diameter of 8.5 cm), each made of 16 AWG wire loops with four or five evenly spaced capacitors. All elements are tuned to 297.2 MHz. (**c**) The coil former has slightly larger dimensions than the brain holder, which slides inside a volume coil (**b**). (**d**) A custom air-tight brain holder was designed to conform to the shape of a whole human brain. The brain holder is an oblate spheroid container (16 × 19 cm) with degassing ports that are used to apply a vacuum suction, thereby reducing air bubbles in the specimen and surrounding fomblin solution.

The coil former (Fig. 2c) consists of two halves and encloses the brain holder. The receive array coil consists of 31 detectors (Fig. 2a), with 15 elements on the top half (diameter = 5.5 cm) and 16 on the bottom half (diameter = 8.5 cm). Coil elements were constructed using 16 AWG wire loops ^17^, each with four or five evenly spaced capacitors (Supplementary Fig. 1). All elements were tuned to 297.2 MHz and matched to a loaded impedance of 75 Ω to minimize preamplifier noise. Preamplifier decoupling was achieved with a cable length of 6 cm. Preamplifiers were placed directly on the coil elements, yielding a substantial reduction in cable losses compared to a previous 30-channel *ex vivo* brain array^18^. The active detuning circuit was formed across the match capacitor using an inductor and PIN diode.

Tuning, matching, and decoupling of neighboring elements was optimized on the bench with a brain sample immersed in periodate-lysine-paraformaldehyde (PLP) solution. Because coil loading varies with the fixative used, the coil must be tuned and matched on the bench using a brain sample with the correct fixative. (For example, testing can be performed with a brain sample immersed in PLP or formalin, but not the regular loading phantom comprised of water and salt). Loops tuned/matched on PLP showed unloaded-to-loaded quality factor ratio (Q-ratio) of Q_UL_/Q_L_ = 210/20 = 10.5, corresponding to an equivalent noise resistance of 11 ohms for the loaded coil (Q = wL/R). By contrast, formalin is a less lossy fixative, giving a coil Q-ratio of Q_UL_/Q_L_ = 210/60 = 3.5, corresponding to an equivalent noise resistance of 4 ohms.

A shielded detunable volume coil (Fig. 2) was built for excitation, with the following parameters and features: band-pass birdcage, diameter 26.7 cm, and an extended length of 32 cm to accommodate brain samples of larger dimensions. For the detuning circuit we used diodes in every leg of the birdcage. These diodes are powered with the high-power chokes, which can withstand high voltage and short duration inversion pulses.

In summary, this coil system incorporates an improved mechanical design, preamps mounted at the coil detectors, and an extended transmit coil design capable of producing high-power pulses.

### 7 Tesla MRI data acquisition

The brain specimen was scanned on a whole-body human 7 Tesla (7T) Siemens Magnetom MRI scanner (Siemens Healthineers, Erlangen, Germany) with the custom-built coil described above. We utilized a GRE sequence^19^ at 100 μm isotropic spatial resolution with the following acquisition parameters: TR = 40 msec, TE = 14.2 msec, bandwidth = 90 Hz/px, FA = 15°, 20°, 25°, 30°. Total scan time for each FA was 25:01:52 [hh:mm:ss], and each FA acquisition generated 1.98 TB of raw k-space data. To improve the signal-to-noise ratio (SNR) and optimise T_1_ modelling, we collected FLASH scans at four FAs: 15°, 20°, 25°, 30° (Fig. 3). Accounting for localizers, quality assurance (QA) scans, and adjustment scans, the total scan time was 100 hours and 8 minutes, and we collected nearly 7.92 TB of raw k-space data.

**Figure 3.**
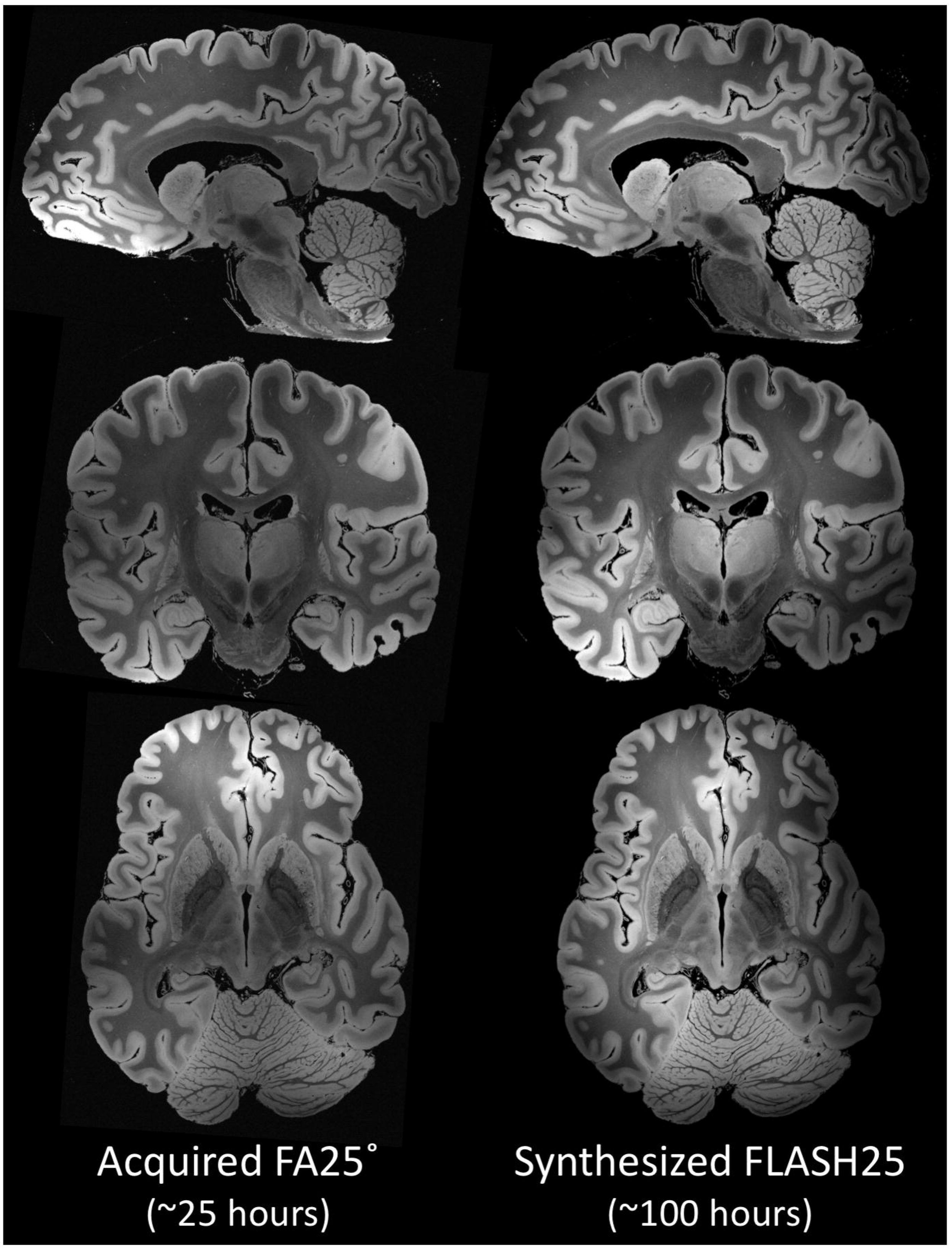
Comparison of FA25° acquisition and synthesized FLASH25 volume. Representative images from the FA25° acquisition (left column) and the synthesized FLASH25 volume (right column) are displayed in the sagittal (top row), coronal (middle row) and axial (bottom row) planes. These images provide a qualitative comparison of the respective signal-to-noise properties of the FA25° acquisition (~25 hours) and the synthesized FLASH25 volume (~100 hours). All images are shown in radiologic convention.

### MRI data reconstruction

The size of the k-space data exceeded the storage capacity of the RAID provided by the scanner image reconstruction computer. The image reconstruction also required more RAM than what was available. We therefore implemented software on the scanner to stream the data directly via TCP/IP to a server on an external computer added to the scanner network, which saved the data as they were received. Because of additional limitations related to the total size of the raw data for any single scan, as dictated by the imager RAID size, we also divided each acquisition into segments. The server on the external computer stored the data as they were acquired, creating date stamps for every k-space segment.

After the scan was completed, the streamed k-space data were transferred to a computational server where we ran custom software to stitch together the segments, reconstruct the images for each channel (via a 3D FFT on each volume per channel^20^), and combine the images derived from the 31 channels via the root-sum-of-squares of the signal magnitudes at each voxel. These signal magnitudes were channel-wise decorrelated using a covariance matrix of the channels’ thermal noise. The output from coil combination was the final acquired image (Data Citation 1; Videos 1, 2 and 3).

### MRI data processing

The acquired data underwent a series of processing steps, culminating in the creation of a T_1_ parameter map and synthesized FLASH volumes (Fig. 3 and Fig. 4; Videos 4, 5, and 6; Data Citations 1 and 2). The volumes were estimated directly from the four FLASH acquisitions using the DESPOT1 algorithm^19,21^ with the program ‘mri_ms_fitparms’ distributed in FreeSurfer (http://surfer.nmr.mgh.harvard.edu) to quantify tissue properties independent of scanner and sequence types. This algorithm fits the tissue parameters (i.e. T1) of the signal equation for the FLASH scan at each voxel using multiple input volumes. The volumes at the originally acquired TRs and flip angles were then regenerated from the parameter maps by evaluating the FLASH signal equation. In principle, a volume with any TR and flip angle combination could be synthesized. These synthesized volumes are created from all the acquired data, and therefore have better SNR than the individually acquired input volumes. We choose to release the 25 degree synthetic volume as it has maximal SNR and the best apparent contrast for cortical and subcortical structures^9^.

**Figure 4.**
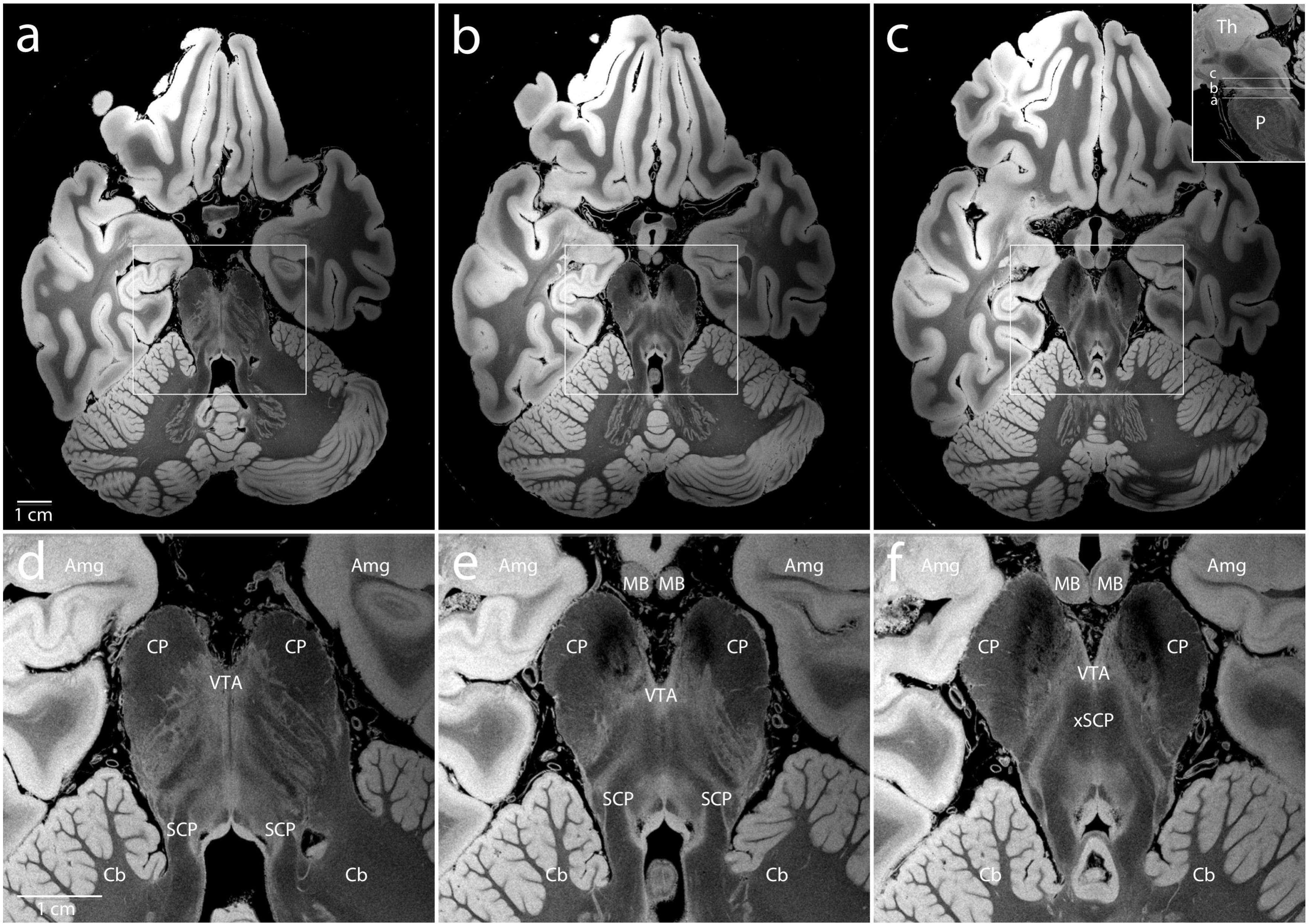
Delineation of subcortical neuroanatomy. Representative axial sections from the synthesized FLASH25 volume are shown at the level of the rostral pons and caudal midbrain (**a-c**, see inset in panel **c**). Zoomed views of the brainstem, medial temporal lobe, and anterior cerebellum (within the white rectangles in **a-c**) are shown in the bottom row (**d-f**). The anatomic detail that can be visualized in this *ex vivo* 100 μm resolution MRI dataset is beyond that which can be seen in typical *in vivo* MRI datasets. All images are shown in radiologic convention. Neuroanatomic abbreviations: Amg = amygdala; Cb = cerebellum; CP = cerebral peduncle; MB = mammillary body; P = pons; SCP = superior cerebellar peduncle; VTA = ventral tegmental area; xSCP = decussation of the superior cerebellar peduncle; Th = thalamus.

Of note, *ex vivo* MRI of the fixed human brain yields a different contrast than *in vivo* MRI, mainly from a shortened T_1_, but also from a decrease in T_2_^*^, both of which are related to formalin fixation^22^. The predominant source of signal contrast in *ex vivo* MRI is likely myelin^23^ and/or iron^24^. Specifically, myelin appears to be a source of T_1_ contrast, while cortical iron appears to be a source of T_2_^*^ contrast^25^.

### Coregistration of the dataset to standard stereotactic space

The dataset was spatially normalized into the MNI ICBM 2009b NLIN ASYM template^26^ (Supplementary Fig. 2a). This template constitutes the newest version of the “MNI space” and is considered a high-resolution version of MNI space because it is available at 0.5 mm isotropic resolution. To combine structural information present on T_1_ and T_2_ versions of the template, we created a joint template using PCA, as previously described^27^. The four synthesized FLASH volumes (FA15, FA20, FA25, and FA30) were downsampled to isotropic voxel-sizes of 0.5 mm for spatial normalization and initially registered into template space in a multispectral approach using Advanced Normalization Tools (ANTs; http://stnava.github.io/ANTs/; ^28^). This multispectral approach simultaneously accounts for intensity data in all four volumes. The initial normalization was performed in four stages (rigid body, affine, whole brain SyN and subcortically focused SyN) as defined in the “effective: low variance + subcortical refine” preset implemented in Lead-DBS software (www.lead-dbs.org; ^29^).

To refine the warp, we introduced fiducial regions of interest (ROI) iteratively using a tool developed for this task (available within Lead-DBS). Specifically, we manually drew line and point fiducial markers in both native and template spaces (Supplementary Fig. 2b). In addition, we manually segmented four structures in native space (subthalamic nucleus, internal and external pallidum and red nucleus). The three types of fiducials (line ROI, spherical ROI and manual segmentations of key structures) were then added as “spectra” in subsequent registration refinements (Supplementary Fig. 2c). Thus, the final registration consisted of a large number of pairings between native and template space (the first four being the actual anatomical volumes, the subsequent ones being manual segmentations and paired helper fiducials). To achieve maximal registration precision, the warp was refined in over 30 iterations with extensive manual expert interaction, each refinement continuing directly from the last saved state. We used linear interpolation to create the normalized files in the data release (Data Citations 1 and 3).

### Code availability

Neuroimaging data were processed using standard processing pipelines (http://surfer.nmr.mgh.harvard.edu/, https://github.com/freesurfer/freesurfer). All code used for registration of volumes into standard stereotactic space are available within the open-source Lead-DBS software (https://github.com/leaddbs/leaddbs). Because registration involved multiple manual user interface steps, no ready-made code is provided, but the process can be readily reproduced with the provided data and software.

## Data Records

The native space FA25° acquisition and synthesized FLASH25 volume are available for download at https://datadryad.org (Data Citation 1). Additional synthesized volumes are available upon request to the corresponding author. Axial, coronal, and sagittal videos of the native space FA25° acquisition (Videos 1, 2, and 3) and synthesized FLASH25 volume (Videos 4, 5, and 6) are also available at the Dryad data repository (Data Citation 1). The synthesized FLASH25 volume is available for interactive, online viewing at https://histopath.nmr.mgh.harvard.edu (Data Citation 2). The normalized FLASH25 volume in standard stereotactic space is available at the Dryad data repository (Data Citation 1) and is hosted on www.lead-dbs.org (preinstalled as part of the LEAD-DBS software package; Data Citation 3).

## Technical Validation

### Coil signal-to-noise ratio (SNR) measurements

The receive coil has a Q_UL_/Q_L_ ratio that ranged from 6 in the top half elements to 8 in the bottom half elements due to larger coil diameter. The S_12_ coupling between neighbouring elements, measured with all other coils active detuned, ranged from −10.9 to −24 dB. All individual elemnts had S_11_ < −20 dB and active detuning of > 30 dB. We evaluated the performance of the transmit coil by examining the B_1_^+^ profile^14^, which shows the efficiency throughout the entire spatial distribution of the brain specimen. The efficiency was greatest in the center of the specimen and fell off gradually towards the edges, as expected for a whole brain specimen at 7T.

We compared the SNR of the 31-channel *ex vivo* array to that of a standard 31-channel 7T head coil and a 64-channel 3T head coil. SNR maps were computed following the method of Kellman & McVeigh^30^. We calibrated the voltage required for 180° pulse using a B_1_^+^ map (estimated with the AFI method)^31^ with an ROI of 3-cm diameter at the center of the brain. We estimated array noise covariance from thermal noise data acquired without RF excitation. The SNR gain with the 31-channel *ex vivo* array was 1.6-fold versus the 31-channel 7T standard coil and 3.3-fold versus the 64-channel 3T head array (Fig. 5). The noise coupling between channels was 11% for the 31-channel *ex vivo* array, a 2-fold improvement relative to our previous array^18^.

**Figure 5.**
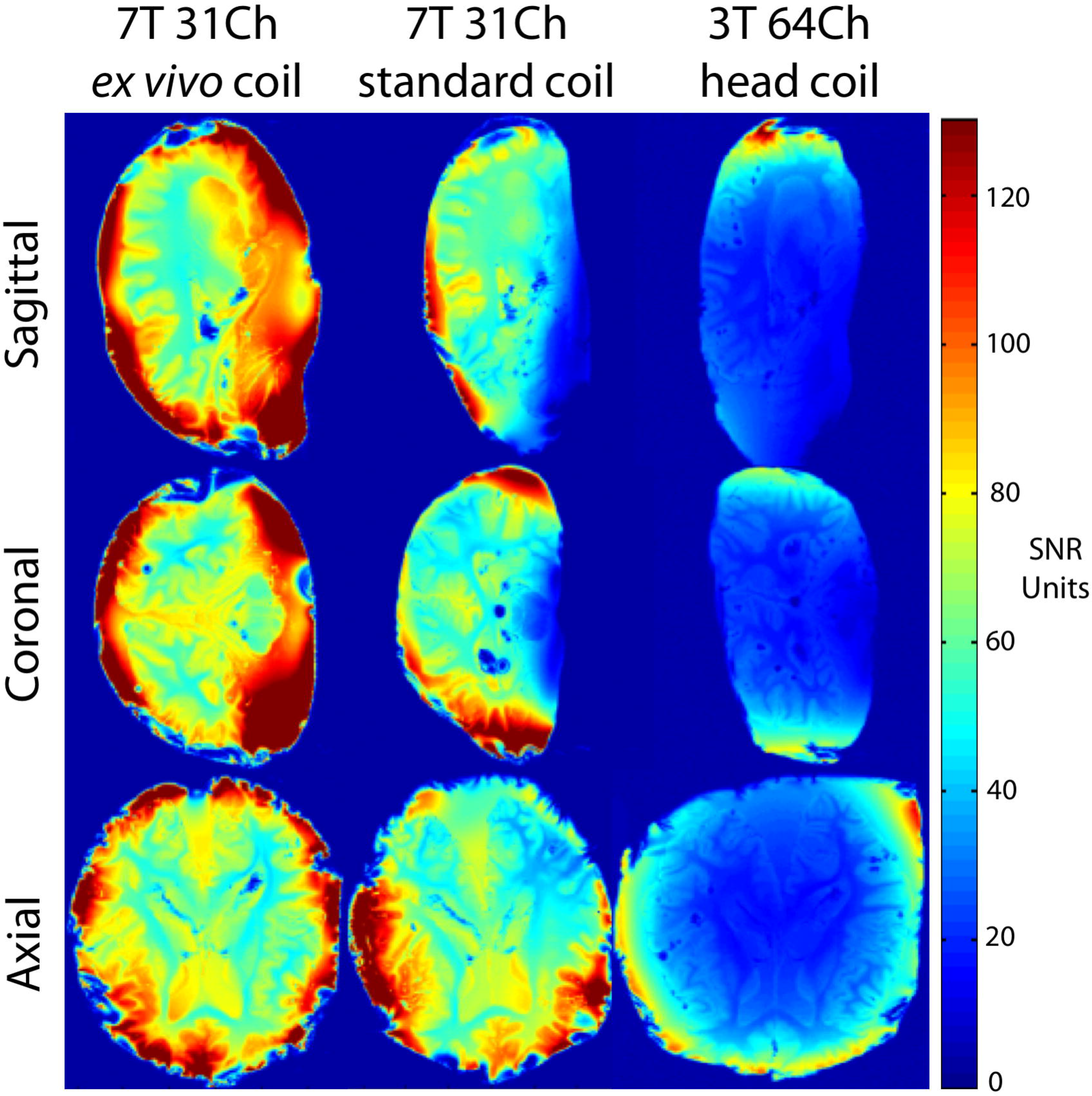
Signal-to-noise ratio (SNR) analysis of coil performance. Representative SNR maps are shown in the sagittal (top row), coronal (middle row) and axial (bottom row) planes for a test brain sample immersed in periodate-lysine-paraformaldehyde. The maps demonstrate an SNR gain of 1.6-fold for the 31-channel 7 Tesla (7T) *ex vivo* coil (left column) compared to the 31-channel 7T standard coil (middle row), and a gain of 3.3-fold compared to the 64-channel 3T head coil (right column). The noise coupling between channels was 11% for the 31-channel *ex vivo* coil array, a 2-fold improvement relative to our previous array^18^.

### Coregistration accuracy

We assessed the neuroanatomic accuracy of the final registration results (i.e. the fit between structures on the normalized FLASH volumes versus the high-resolution MNI template) by visual inspection using a tool specifically designed for this task (implemented in Lead-DBS). An example of this visual inspection assessment for the subthalamic nucleus and globus pallidus interna is provided in Supplementary Fig. 3. The final maps are stored in NIfTI and mgz files in isotropic 150 μm resolution (Data Citation 1). The normalized FLASH25 volume is additionally distributed pre-installed within Lead-DBS software and can be selected for visualization in the 3D viewer (Data Citation 3). Fig. 6 shows an example in synopsis with deep brain stimulation electrode reconstructions in a hypothetical patient being treated for Parkinson’s Disease.

**Figure 6.**
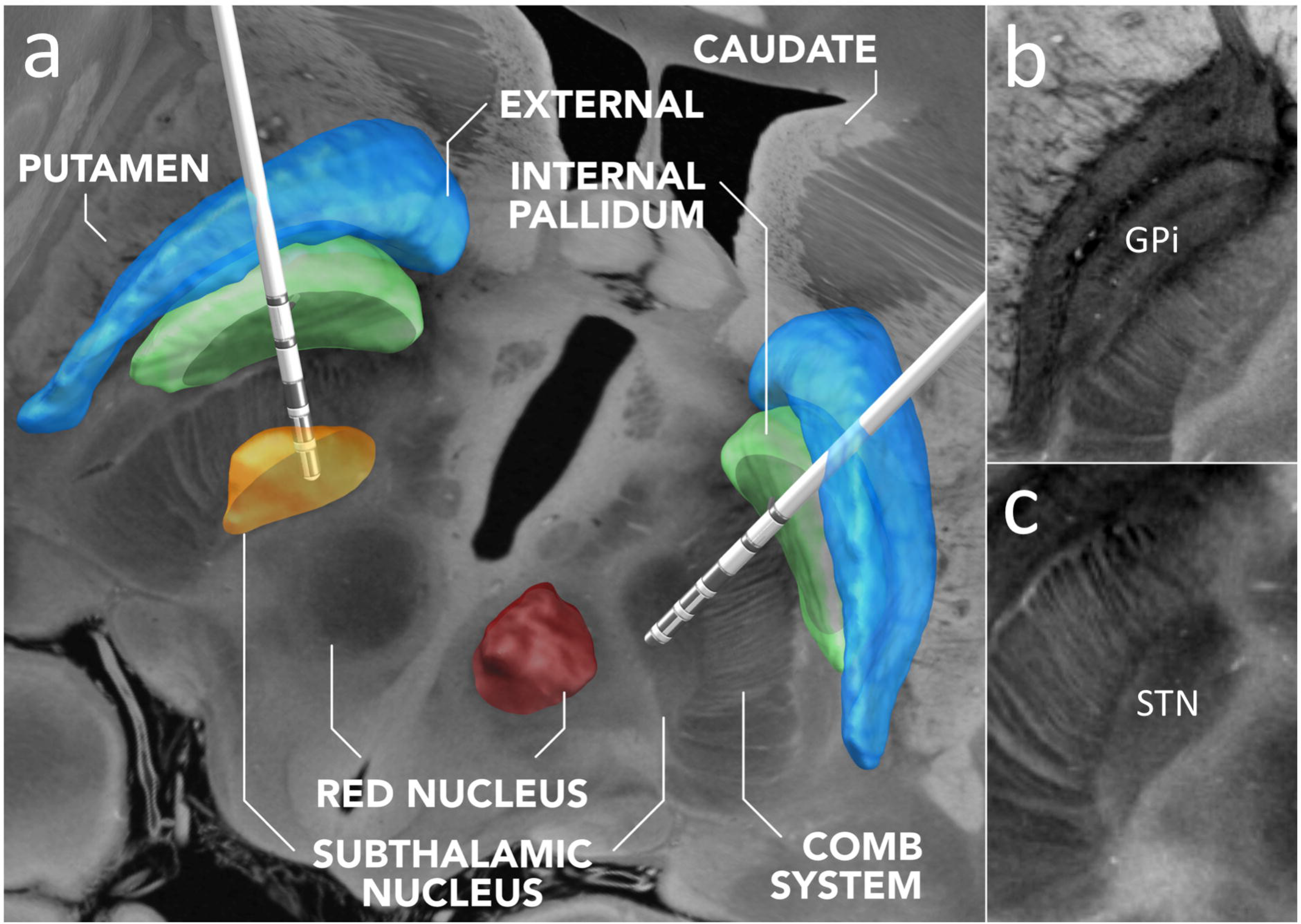
Normalization of the ex vivo MRI dataset into standard stereotactic space and integration into the Lead-DBS software platform. **(a)** Exemplary use-case of the normalized FLASH25 volume in deep brain stimulation (DBS). DBS electrodes are visualized for a hypothetical patient using Lead-DBS software (https://www.lead-dbs.org)^29^. An axial image from the normalized scan, at the level of the rostral midbrain, is shown as a backdrop, with 3D-structures defined by the DISTAL atlas^32^ (right subthalamic and left red nucleus hidden for optimal visualization of the underlying anatomy). Panels (**b**) and (**c**) show zoomed views of key DBS target regions: the left globus pallidus interna (GPi in **b**) and the subthalamic nucleus (STN in **c**). The images in (**b**) and (**c**) are shown in radiologic convention.

## Acknowledgements

We thank Michelle Siciliano and Terrence Ott for assistance in obtaining and processing the brain specimen. We thank Simon Sigalovsky for assistance with coil construction, and Gunjan Madan for assistance with coil testing and evaluation. We thank L. Daniel Bridgers for constructing the brain container and coil array housing. We thank Andrew Hoopes for assistance with creation of visual media. This work was supported by the National Institutes of Health (NIH) National Institute for Neurological Disorders and Stroke (K23-NS094538, R01-NS052585, R21-NS072652, R01-NS070963, R01-NS083534, U01-NS086625), National Institute for Biomedical Imaging and Bioengineering (P41-EB015896, R01-EB006758, R21-EB018907, R01-EB019956, R01-EB023281, R00-EB021349), National Institute on Aging (R01-AG057672, R01-AG022381, R01-AG008122, R01-AG016495, R01-AG008122, U01-AG006781, R21-AG046657, P41-RR014075, P50-AG005136), National Center for Alternative Medicine (RC1-AT005728), Eunice Kennedy Shriver National Institute of Child Health and Human Development (K01-HD074651, R01-HD071664, R00-HD074649), and the Centers for Disease Control and Prevention (R49-CE001171). Support for this research was also provided by the BRAIN Initiative Cell Census Network (U01-MH117023) and the NIH Blueprint for Neuroscience Research (U01-MH093765) as part of the multi-institutional Human Connectome Project. This research utilized resources provided by the National Center for Research Resources (U24-RR021382) and by NIH shared instrumentation grants S10-RR023401, S10-RR019307, and S10-RR023043. Additional support was provided by the James S. McDonnell Foundation, Rappaport Foundation, the Tiny Blue Dot Foundation as well as the German Research Foundation (Emmy Noether Grant 410169619).

## Author contributions

B.L.E. designed the study, analyzed the data, and prepared the manuscript.

A.M. built the coil, acquired and analyzed the data, and contributed to the manuscript.

A.H. created the warp from native space to standard stereotactic space, performed the coregistration for Lead-DBS implementation, and contributed to the manuscript.

J.R.P. designed the study, acquired and analyzed the data, and contributed to the manuscript.

T.W. designed the study, acquired the data, and contributed to the manuscript.

M.D.T. acquired and analyzed the data, and contributed to the manuscript.

J.A. designed the study, acquired and analyzed the data, and contributed to the manuscript.

J.P.S. advised on the building and testing of the coil, and contributed to the manuscript.

B.R.D. analyzed the data and contributed to the manuscript.

A.S. acquired and analyzed the data, and contributed to the manuscript.

L.S.T. processed and analyzed the data, and contributed to the manuscript.

R.D.F. performed the pathological assessment and contributed to the manuscript.

L.L.W. supervised the building of the coil and contributed to the manuscript.

B.F. supervised and designed the study, analyzed the data, and contributed to the manuscript.

A.v.d.K. supervised and designed the study, acquired and analyzed the data, and contributed to the manuscript.

## Additional Information

### Competing interests

None of the authors has a conflicting financial interest. Dr. Fischl and Mr. Tirrell have financial interest in CorticoMetrics, a company whose medical pursuits focus on brain imaging and measurement technologies. Their interests were reviewed and are managed by Massachusetts General Hospital and Partners HealthCare in accordance with their conflict of interest policies.

## Videos

**Video 1. Axial images from the FA25° acquisition.** These images were acquired in ~25 hours of scan time. The images are shown in radiologic convention.

**Video 2. Coronal images from the FA25° acquisition.** These images were acquired in ~25 hours of scan time. The images are shown in radiologic convention.

**Video 3. Sagittal images from the FA25° acquisition.** These images were acquired in ~25 hours of scan time.

**Video 4. Axial images from the synthesized FLASH25 volume.** These images were acquired in ~100 hours of scan time. The images are shown in radiologic convention.

**Video 5. Coronal images from the synthesized FLASH25 volume.** These images were acquired in ~100 hours of scan time. The images are shown in radiologic convention.

**Video 6. Sagittal images from the synthesized FLASH25 volume.** These images were acquired in ~100 hours of scan time.

## Notes

#### Summary of Updates

Corrected author list. Added source of funding.

https://datadryad.org

https://www.youtube.com/playlist?list=PLlL7CEMX5bxzl1TqkJRK3pMV4Ax6By4bN

https://histopath.nmr.mgh.harvard.edu

https://www.lead-dbs.org

## REFERENCES

1 Augustinack, J. C., van der Kouwe, A. J. & Fischl, B. Medial temporal cortices in ex vivo magnetic resonance imaging. J Comp Neurol 521, 4177–4188, doi:10.1002/cne.23432 (2013).

2 Edlow, B. L. et al. Neuroanatomic connectivity of the human ascending arousal system critical to consciousness and its disorders. J Neuropathol Exp Neurol 71, 531–546, doi:10.1097/NEN.0b013e3182588293 (2012).

3 McNab, J. A. et al. The Human Connectome Project and beyond: Initial applications of 300mT/m gradients. Neuroimage 80, 234–245, doi:10.1016/j.neuroimage.2013.05.074 (2013).

4 Roebroeck, A., Miller, K. L. & Aggarwal, M. Ex vivo diffusion MRI of the human brain: Technical challenges and recent advances. NMR Biomed 32, e3941, doi:10.1002/nbm.3941 (2019).

5 Aggarwal, M. et al. Feasibility of creating a high-resolution 3D diffusion tensor imaging based atlas of the human brainstem: a case study at 11.7 T. Neuroimage 74, 117–127, doi:10.1016/j.neuroimage.2013.01.061 (2013).

6 Lusebrink, F., Sciarra, A., Mattern, H., Yakupov, R. & Speck, O. T1-weighted in vivo human whole brain MRI dataset with an ultrahigh isotropic resolution of 250 mum. Sci Data 4, 170032, doi:10.1038/sdata.2017.32 (2017).

7 Horn, A. et al. Teaching NeuroImages: In vivo visualization of Edinger comb and Wilson pencils. Neurology 92, e1663–e1664, doi:10.1212/WNL.0000000000007252 (2019).

8 Takahashi, E., Song, J. W., Folkerth, R. D., Grant, P. E. & Schmahmann, J. D. Detection of postmortem human cerebellar cortex and white matter pathways using high angular resolution diffusion tractography: a feasibility study. Neuroimage 68, 105–111, doi:10.1016/j.neuroimage.2012.11.042 (2013).

9 Augustinack, J. C. et al. Detection of entorhinal layer II using 7Tesla [corrected] magnetic resonance imaging. Ann Neurol 57, 489–494, doi:10.1002/ana.20426 (2005).

10 Gangolli, M. et al. Quantitative validation of a nonlinear histology-MRI coregistration method using generalized Q-sampling imaging in complex human cortical white matter. Neuroimage 153, 152–167, doi:10.1016/j.neuroimage.2017.03.059 (2017).

11 van Veluw, S. J. et al. Microbleed and microinfarct detection in amyloid angiopathy: a high-resolution MRI-histopathology study. Brain 139, 3151–3162, doi:10.1093/brain/aww229 (2016).

12 Sengupta, S. et al. High resolution anatomical and quantitative MRI of the entire human occipital lobe ex vivo at 9.4T. Neuroimage 168, 162–171, doi:10.1016/j.neuroimage.2017.03.039 (2018).

13 van der Kouwe, A. et al. High Resolution Structural and Diffusion MRI of Ex Vivo Human Motor Cortex. Organization for Human Brain Mapping (2011).

14 Mareyam, A. et al. Array coil and sample preparation and support system for whole brain ex vivo imaging at 100 μm. ISMRM. #3130 (2015).

15 Edlow, B. L. et al. Multimodal Characterization of the Late Effects of Traumatic Brain Injury: A Methodological Overview of the Late Effects of Traumatic Brain Injury Project. J Neurotrauma 35, 1604–1619, doi:10.1089/neu.2017.5457 (2018).

16 Bridgers, L. D. Design and Manufacture of an Ultra-High Field Ex Vivo Coil Assembly, Massachusetts Institute of Technology, (2012).

17 Keil, B. et al. Size-optimized 32-channel brain arrays for 3 T pediatric imaging. Magn Reson Med 66, 1777–1787, doi:10.1002/mrm.22961 (2011).

18 Mareyam, A., Polimeni, J. R., Alagappan, V., Fischl, B. & Wald, L. L. A 30 channel receive-only 7T array for ex vivo brain hemisphere imaging. ISMRM. #106 (2009).

19 Fischl, B. et al. Sequence-independent segmentation of magnetic resonance images. Neuroimage 23 Suppl 1, S69–84, doi:10.1016/j.neuroimage.2004.07.016 (2004).

20 Frigo, M. & Johnson, S. G. The Design and Implementation of FFTW3. Proceedings of the IEEE 93, 216–231 (2005).

21 Deoni, S. C., Peters, T. M. & Rutt, B. K. High-resolution T1 and T2 mapping of the brain in a clinically acceptable time with DESPOT1 and DESPOT2. Magn Reson Med 53, 237–241, doi:10.1002/mrm.20314 (2005).

22 Tovi, M. & Ericsson, A. Measurements of T1 and T2 over time in formalin-fixed human whole-brain specimens. Acta Radiol 33, 400–404 (1992).

23 Eickhoff, S. et al. High-resolution MRI reflects myeloarchitecture and cytoarchitecture of human cerebral cortex. Hum Brain Mapp 24, 206–215, doi:10.1002/hbm.20082 (2005).

24 Fukunaga, M. et al. Layer-specific variation of iron content in cerebral cortex as a source of MRI contrast. Proc Natl Acad Sci U S A 107, 3834–3839, doi:10.1073/pnas.0911177107 (2010).

25 Stuber, C. et al. Myelin and iron concentration in the human brain: a quantitative study of MRI contrast. Neuroimage 93 Pt 1, 95–106, doi:10.1016/j.neuroimage.2014.02.026 (2014).

26 Fonov, V. S., Evans, A. C., McKinstry, R. C., Almli, C. R. & Collins, D. Unbiased nonlinear average age-appropriate brain templates from birth to adulthood. Neuroimage 47, S102 (2009).

27 Horn, A. PCA MNI 2009b NLIN template. (2017). <doi:10.6084/m9.figshare.4644472.v2>.

28 Avants, B. B., Epstein, C. L., Grossman, M. & Gee, J. C. Symmetric diffeomorphic image registration with cross-correlation: evaluating automated labeling of elderly and neurodegenerative brain. Medical image analysis 12, 26–41, doi:10.1016/j.media.2007.06.004 (2008).

29 Horn, A. et al. Lead-DBS v2: Towards a comprehensive pipeline for deep brain stimulation imaging. Neuroimage 184, 293–316, doi:10.1016/j.neuroimage.2018.08.068 (2019).

30 Kellman, P. & McVeigh, E. R. Image reconstruction in SNR units: a general method for SNR measurement. Magn Reson Med 54, 1439–1447, doi:10.1002/mrm.20713 (2005).

31 Yarnykh, V. L. Actual flip-angle imaging in the pulsed steady state: a method for rapid three-dimensional mapping of the transmitted radiofrequency field. Magn Reson Med 57, 192–200, doi:10.1002/mrm.21120 (2007).

32 Ewert, S. et al. Toward defining deep brain stimulation targets in MNI space: A subcortical atlas based on multimodal MRI, histology and structural connectivity. Neuroimage 170, 271–282, doi:10.1016/j.neuroimage.2017.05.015 (2018).

## Data Citations

1. Edlow, B.L., Mareyam, A., Horn, A., Polimeni, J.R., Witzel, T., Tisdall, M.D., Augustinack, J., Stockmann, J.P., Diamond B.R., Stevens’ A., Tirrell, L., Folkerth, R.D., Wald, L.L., Fischl, B. & Kouwe, A.v.d. Dryad Digital Repository. https://datadryad.org. doi:10.5061/dryad.119f80q (2019).

2. Edlow, B.L., Mareyam, A., Horn, A., Polimeni, J.R., Witzel, T., Tisdall, M.D., Augustinack, J., Stockmann, J.P., Diamond B.R., Stevens’ A., Tirrell, L., Folkerth, R.D., Wald, L.L., Fischl, B. & Kouwe, A.v.d. Biolucida https://histopath.nmr.mgh.harvard.edu (2019).

3. Edlow, B.L., Mareyam, A., Horn, A., Polimeni, J.R., Witzel, T., Tisdall, M.D., Augustinack, J., Stockmann, J.P., Diamond B.R., Stevens’ A., Tirrell, L., Folkerth, R.D., Wald, L.L., Fischl, B. & Kouwe, A.v.d. Lead-DBS https://www.lead-dbs.org (2019).

