## Supplementary Material for "7 Tesla MRI of the *ex vivo* human brain at 100 micron resolution"

* co-senior authors

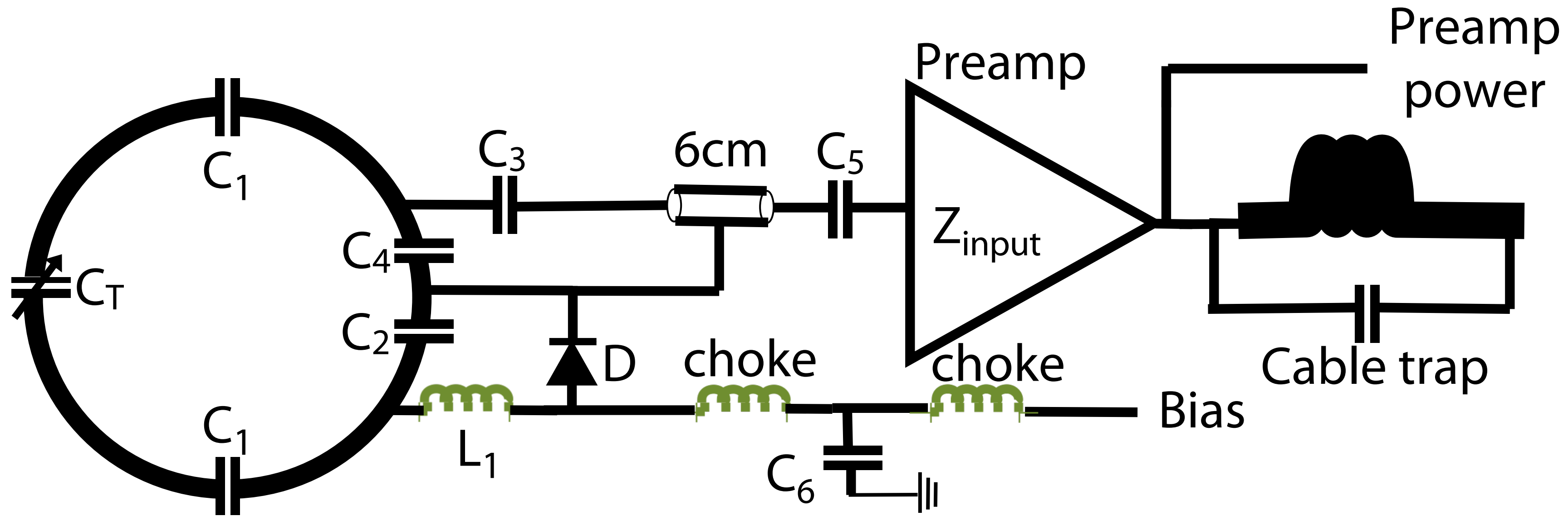


**Supplementary Figure 1. Circuit diagram of each coil element.** All elements were constructed using 16 AWG wire loops[1], each with four or five evenly spaced capacitors. All elements were tuned to 297.2 MHz and matched to a loaded impedance of 75 Ω to minimize preamplifier noise. Preamplifier decoupling was achieved with a cable length of 6 cm. Preamplifiers were placed directly on the coil elements, yielding a substantial reduction in cable losses compared to a previous 30-channel *ex vivo* brain array[2].

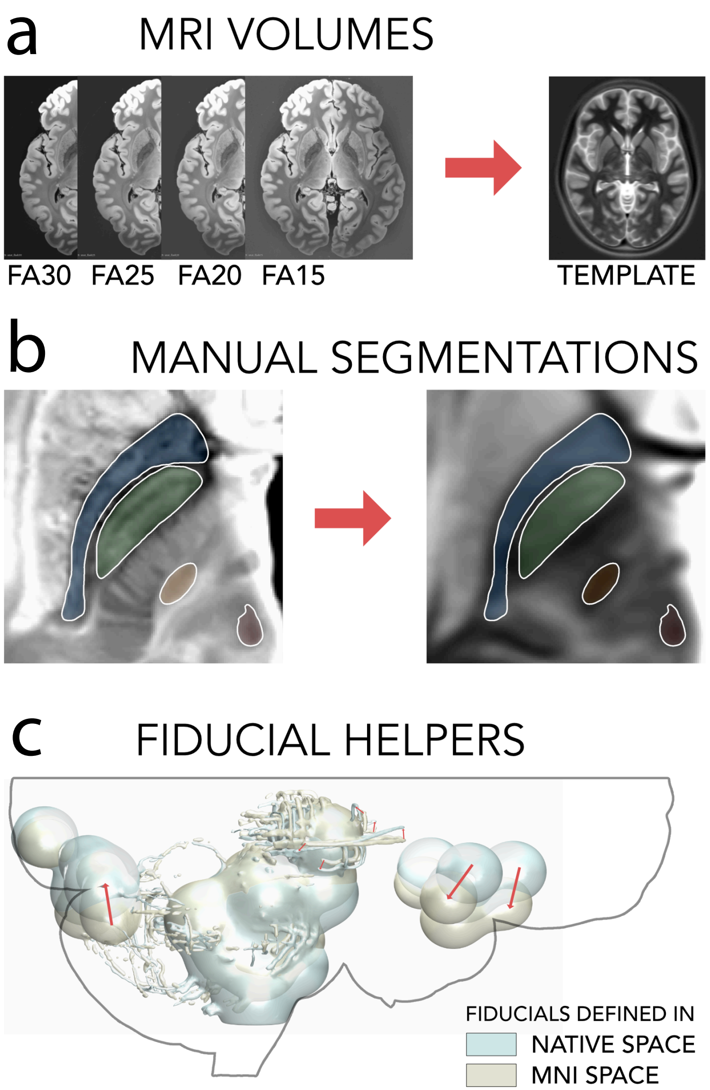


**Supplementary Figure 2. Coregistration to standard stereotactic space.** (**a**) We performed a multispectral warp between FA15-30 volumes and the PCA template of the ICBM 2009b NLIN ASYM space[3,4], which built the basis of the iterative registration process. (**b**) In addition, we manually segmented key subcortical structures – the subthalamic nucleus (orange), red nucleus (red), internal (green) and external (blue) pallidum – in native space and paired them with corresponding structures in MNI space, as defined by the DISTAL atlas [5]. (**c**) Additionally, we manually introduced line and smoothed sphere fiducial markers into native and MNI spaces, and we paired them in the registration process. Registration was performed in ~30 iterations, and fiducial corrections (shown in (**c**)) were introduced iteratively to further optimize fit.

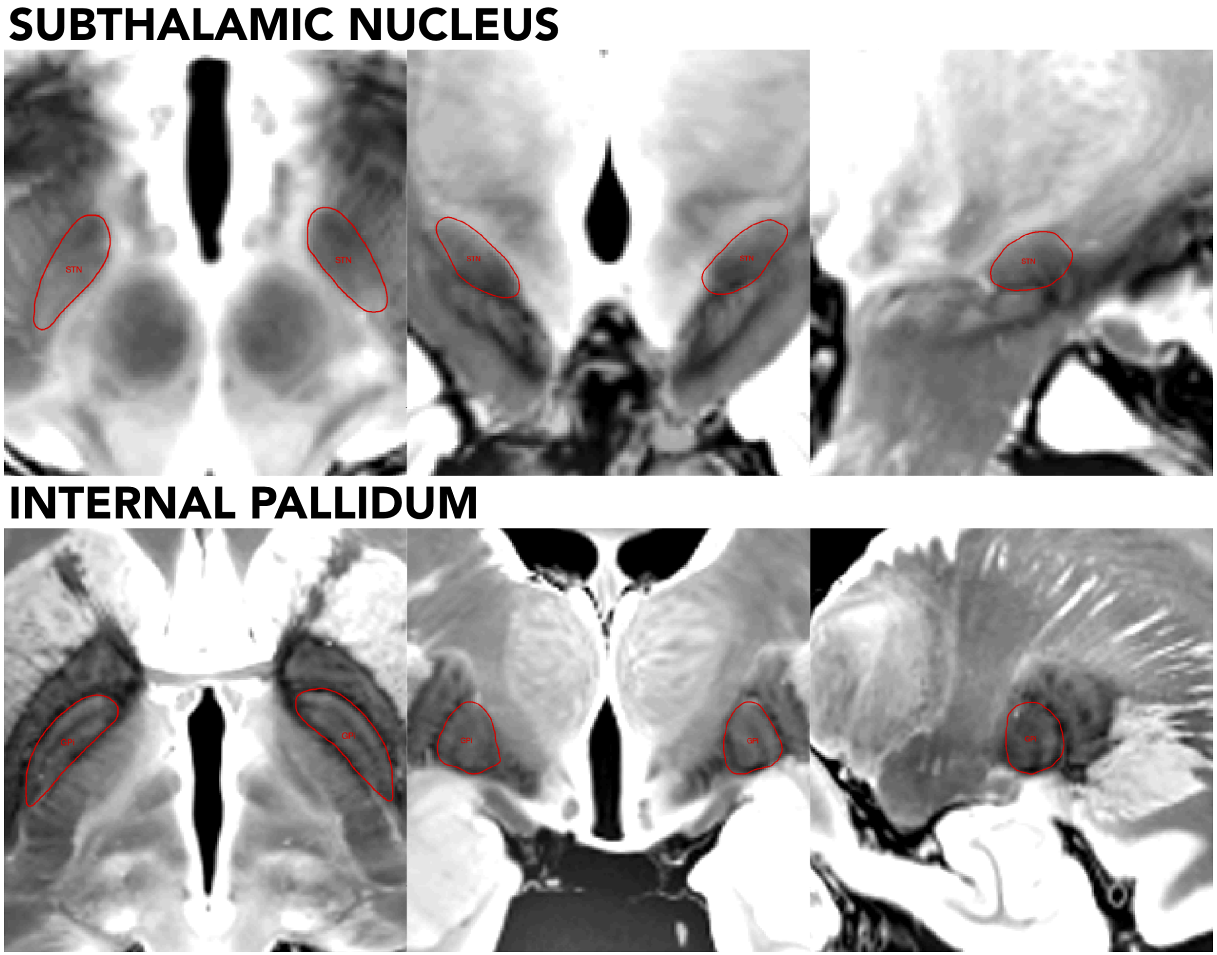


**Supplementary Figure 3. Neuroanatomic accuracy of the coregistration to stereotactic space.** We show the registration fit for subcortical structures that are clinically relevant to deep brain stimulation: subthalamic nucleus (STN) and internal pallidum (GPi). Red lines show the boundaries of these structures defined by the DISTAL atlas[5]. The normalized FA25 volume is shown in the background (left column = axial, middle column = coronal, right column = sagittal views).

### Supplementary References


1 Keil, B. *et al.* Size-optimized 32-channel brain arrays for 3 T pediatric imaging. *Magn Reson Med* **66**, 1777-1787, doi:10.1002/mrm.22961 (2011).

2 Mareyam, A., Polimeni, J. R., Alagappan, V., Fischl, B. & Wald, L. L. A 30 channel receive-only 7T array for ex vivo brain hemisphere imaging. *ISMRM. #106* (2009).

3 Horn, A. PCA MNI 2009b NLIN template. doi:10.6084/m9.figshare.4644472.v2 (2017).

4 Fonov, V. S., Evans, A. C., McKinstry, R. C., Almli, C. R. & Collins, D. Unbiased nonlinear average age-appropriate brain templates from birth to adulthood. *Neuroimage* **47**, S102 (2009).

5 Ewert, S. *et al.* Toward defining deep brain stimulation targets in MNI space: A subcortical atlas based on multimodal MRI, histology and structural connectivity. *Neuroimage* **170**, 271-282, doi:10.1016/j.neuroimage.2017.05.015 (2018).
